# Methods for detecting “missing” dimensions in genetic covariance matrices

**DOI:** 10.1101/2024.07.29.605681

**Authors:** Ned A. Dochtermann

## Abstract

Blows and Hoffmann (2005) and others have suggested that low levels of genetic variation in some dimensions of an additive genetic variance-covariance matrix (**G**) will be detectable as eigenvalues approaching zero. These dimensions may be indicative of trade-offs (Blows and Hoffmann 2005) or, potentially, of “holes” in fitness landscapes (Dochtermann et al. 2023).

However, because the estimation of **G** typically constrains variances to be positive, and matrices to therefore be positive definite, “approaching zero” is challenging to statistically define. It is, therefore, not currently clear how to statistically identify dimensions with effectively zero variances. Being able to identify dimensions of **G** with variances approaching zero would improve our ability to understand trade-offs which, typically and inappropriately, focus on the signs of bivariate correlations (Houle 1991). Viable approaches would also allow better understanding of the structure of **G** and the evolutionary processes that have shaped **G** across taxa.

Mezey and Houle (2005) and Kirkpatrick and Lofsvold (1992) used matrix rank to determine if there were dimensions lacking variation. Rank is the number of eigenvalues for a matrix that are greater than zero. If a **G**’s rank is less than its dimensionality, this would be evidence that there are dimensions without variation. However, most contemporary analysis procedures force estimates of **G** to be positive semi-definite. Consequently, estimated **G**s will not have eigenvalues of 0 or negative. Therefore, the estimated eigenvalues on their own will not allow the identification of absolute constraints (*sensu* (Houle 2001)) or more generally dimensions lacking variation. One alternative would be to compare the eigenvalues of observed matrices to those of random matrices and, if there were dimensions with less variation than expected by chance, this comparison could be used to identify dimensions in which evolution is constrained.

If trade-offs or landscape holes result in dimensions of **G** that have minimal variance, we would expect that the associated eigenvalues exhibit less variation than expected by chance. Here, I tested whether the distribution of eigenvalues from randomly generated correlated matrices can be used for such testing.

## Methods

I generated random correlation matrices using the LKJ method (Lewandowski et al. 2009) for correlation matrices of various dimensionalities (3, 5, 10, 15, and 20) with η = 1. I then converted these correlation matrices to covariances matrices based on variances that were generated as random deviates pulled from a gamma distribution with shape, *k*, set to 10 and scale, θ, set to 1. This produces large maximum correlations but constraints on permissible matrices result in moderate to small average correlations (Figure 1A). This also produces a wide distribution of variances (Figure 1B). I generated 5000 matrices at each dimensionality.

**Figure 1.**
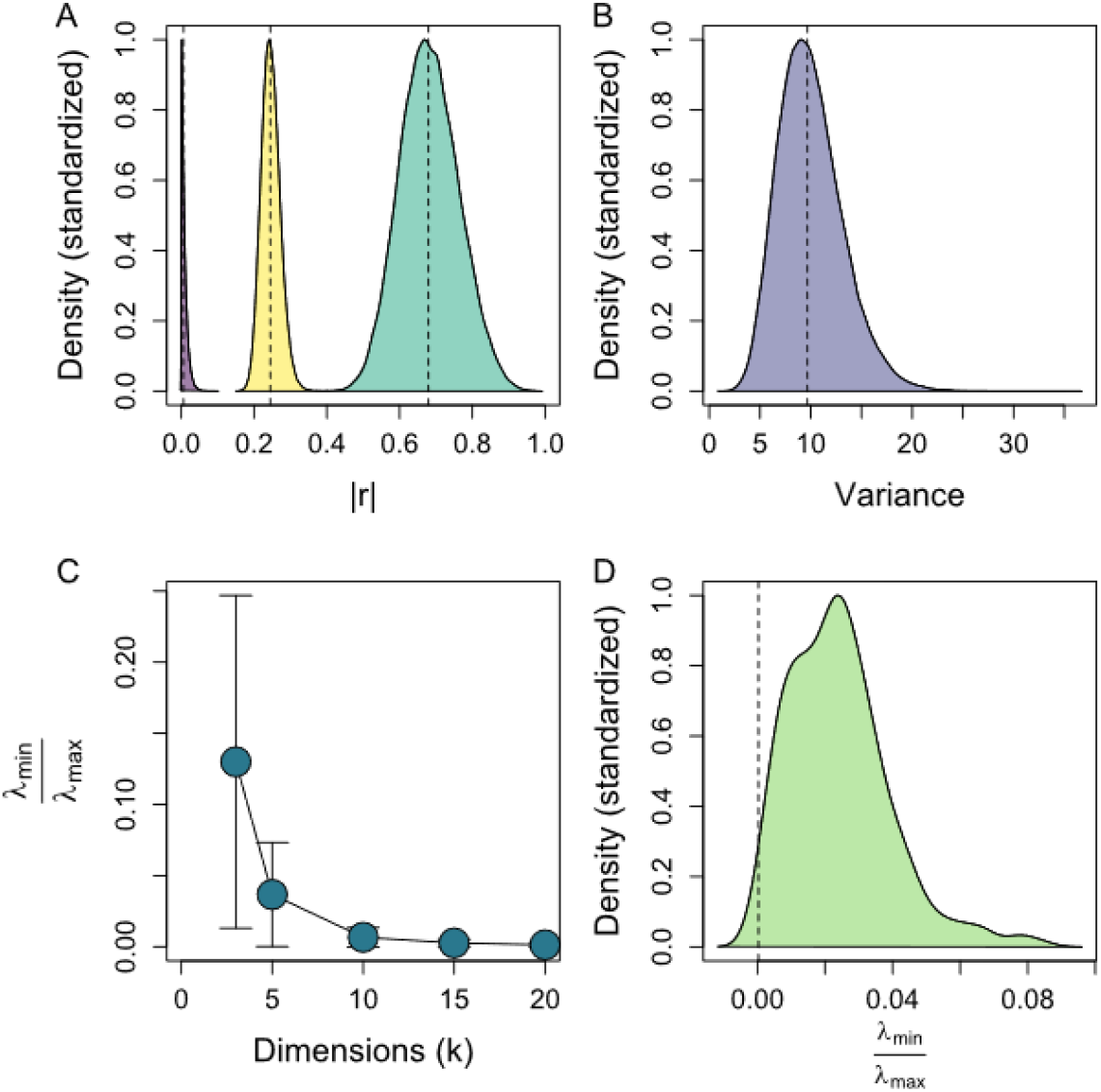
A. Correlations (absolute) produced using the LKJ method as used in our simulations (k = 10, eta = 1, samples = 1e5). The average minimum correlation from LKJ generated matrices was 0.01 (purple). The average maximum correlation was 0.68 (teal). The average mean correlation was 0.25 (yellow). B. Distribution of variances used to convert correlation matrices to covariance matrices. Variances were drawn from a gamma distribution (shape - 10, scale =1). C. Mean ± SD of 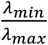 by dimensionality. D. Distribution of 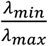 for simulated populations that evolved on Gaussian landscapes via truncation selection.

I then examined the magnitude of each matrix’s minimum eigenvalue relative to the maximum eigenvalue, i.e. 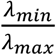 (this is the inverse of condition). This metric was chosen as an eigenvalue “approaching zero” can be defined as containing a small amount of variation relative to the dimension with the most variation. Mathematically, this defines a matrix with eigenvalues “approaching zero” as being an ill-conditioned matrix. The distribution of 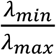 might then provide random distributions and critical cut-off values against which observed values can be compared.

Finally, following the individual variance components simulation procedure described in Dochtermann et al. (2023), I estimated 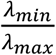 for **G** from 250 populations that evolved on Gaussian fitness landscapes. Selection was imposed as truncation selection with selection set such that only the top 10% of the population was used for the subsequent generation. Populations contained 7500 individuals, individuals had 10 traits, the starting genetic correlations between traits were 0, and populations evolved for 250 generations. The distribution of 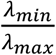 for these populations were compared to cut-off values determined for random correlations.

## Results

For random covariance matrices, dimensions with the least variation typically contain almost *no* variation. When matrices have a dimensionality of 5 or greater, this dimension captures 4% or less of the variation captured in the dominant dimension. Across dimensionalities, the median 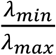 was 0.005 and the maximum was 0.82 (Figure 1C). This maximum was for a 3×3 matrix and 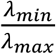 dramatically declined with increasing dimensionality (Figure 1C). Median 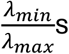 s by dimensionalities were: 0.096, 0.025, 0.0047, 0.0019, and 0.00094 respectively.

Critical cut-off values for indicating deviation from random expectations can be defined based on arbitrary thresholds, for example, a 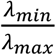 that occurs in less than 5% of random matrices. For random covariance matrices of 3, 5, 10, 15, and 20 dimensions and based on 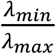 these cut-offs would be: 7.5e^-3^, 2e^-3^, 3.1e^-4^, 1.4e^-4^, and 7.0e^-5^.

Simulated populations exhibited 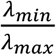 values with a median of 0.023. The distribution of these values largely exceeded those of random matrices (Figure 1D). 94.8% of values were *greater* than the cut-off expected by chance.

## Discussion

Identifying dimensions of **G** matrices with effectively no variation would aid in determining whether constraints on evolutionary responses exist in particular dimensions (Blows and Hoffmann 2005). Dimensions of **G** with no variation might similarly indicate “holes” in fitness landscapes (Dochtermann et al. 2023).

Here, I generated random correlation matrices and analyzed the smallest eigenvalues of these matrices. As dimensionality increased, these smallest eigenvalues approached zero simply due to the constraints imposed by the mathematical requirements of permissible correlation matrices (Figure 1C). Unfortunately, this analysis also demonstrated that these smallest eigenvalues overlapped those of simulated populations that evolved on Gaussian landscapes (Figure 1D). This suggests that random correlation matrices cannot be used to set thresholds for identifying dimensions with minimal variation.

An expansion of the approach described here would be to calculate alternative matrix metrics. Other metrics, for example autonomy or evolvability (Hansen and Houle 2008) may perform better and are directly grounded in evolutionary theory. However, most such metrics rely on the distribution of eigenvalues, as is 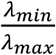 as I used here, so it is not immediately clear that any alternative would allow testing for dimensions that are lacking variation.

Identifying dimensions in multivariate space where the variation present approaches zero would more properly test for trade-offs and constraints than existing approaches. For example, the sign of correlations is not indicative of trade-offs (Houle 1991). Identifying these dimensions may also allow for testing whether “holes” in landscapes are reflected in a population’s **G** (Dochtermann et al. 2023). Unfortunately, based on this preliminary analysis, there are not clear approaches for doing this testing that are currently available to researchers.

